# AAV9-MCT8 delivery at juvenile stage ameliorates neurological and behavioral deficits in an Allan-Herndon-Dudley Syndrome mouse model

**DOI:** 10.1101/2021.10.31.466634

**Authors:** Xiao-Hui Liao, Pablo Avalos, Oksana Shelest, Raz Ofan, Michael Shilo, Catherine Bresee, Shibi Likhite, Jean-Philippe Vit, Heike Heuer, Brian Kaspar, Kathrin Meyer, Alexandra M. Dumitrescu, Samuel Refetoff, Clive N. Svendsen, Gad D. Vatine

## Abstract

Allan-Herndon-Dudley syndrome (AHDS) is a severe X-linked intellectual and psychomotor disability disorder accompanied by abnormal thyroid hormone (TH) levels. AHDS is caused by inactivating mutations in the monocarboxylate transporter 8 (MCT8), a specific TH transporter widely expressed in the central nervous system. *MCT8* gene mutations cause impaired transport of TH across brain barriers, leading to insufficient neural TH supply. There is currently no successful therapy for the neurological symptoms. AAV9-based gene therapy is a promising approach to treat monogenic neurological disorders. Here, the potential of this approach was tested in the well-established double knockout (dKO) Mct8^-/y^;Oatp1c1^-/-^ mouse model of AHDS, which displays disease-relevant neurological and TH phenotypes. Systemic intravenous delivery of AAV9-MCT8 at a juvenile stage led to improved locomotor and cognitive function, as well as rescue of T_3_-brain content and T_3_-related gene expression. This preclinical study indicates that this gene therapy may improve the neurological symptoms of AHDS patients.

## Introduction

Thyroid hormones (THs) are essential for the development and metabolic homeostasis of most organs and tissues^1^. The major form of TH released in the blood from the thyroid gland is thyroxine (T_4_), which acts as a prohormone. T_4_ conversion to the more potent, active hormone triiodothyronine (T_3_) or to the inactive form reverse T_3_ (rT_3_) takes place intracellularly by iodothyronine deiodinases enzymes^2^. The main mechanism of T_3_ action is achieved through binding to specific nuclear receptors, which in turn operate as regulators of gene transcription^3^. Since TH metabolism and action are intracellular events, they require the presence of TH specific transporters mediating cellular TH uptake and efflux^4^. The *solute carrier family 16, member 2* (*SLC16A2*) gene, located on the X-chromosome, encodes for the monocarboxylate transporter 8 (MCT8) protein^5^. MCT8 is well conserved throughout vertebrate evolution and is widely expressed in the body and central nervous system (CNS)^6^. A key function of MCT8 is to facilitate TH transport across plasma membranes^5^.

Inactivating *MCT8* gene mutations in males cause a severe form of psychomotor disability^7–9^, clinically described by Allan Herndon Dudley and now termed (AHDS)^10^. Patients exhibit neurological impairments including severe intellectual disability, truncal hypotonia, dystonia and movement disorders. MCT8-deficiency also causes a TH phenotype, including elevated serum T_3_ levels, low rT_3_ and T_4_ with normal or slightly elevated TSH resulting in markedly elevated free T_3_/T_4_ and T_3_/rT_3_ ratios^11^.

Two independently generated *Mct8*-KO mouse models^12,13^ closely recapitulated the TH phenotype observed in patients with AHDS, but did not display expected neurological or behavioral phenotypes. This was due to a milder TH deprivation in mouse brains owing to a T_4_-specific transporter not present in the human blood brain barrier (BBB). Specifically, the Organic anion-transporting polypeptide 1c1 (Oatp1c1), encoded by the *slco1c1* gene, was identified in mice, but not human, brain capillaries^14–16^. Mice with a double knockout (dKO) of Mct8 and Oatp1c1 displayed disease-relevant phenotypes including an impaired TH transport into the CNS and consequently a significantly decreased number of cortical parvalbumin-positive GABAergic interneurons, reduced myelination and pronounced locomotor abnormalities^17^. These results indicated that in mice Mct8 (together with Oatp1c1) plays a crucial role in the transport of THs into the CNS and, importantly, provided a robust disease model for human MCT8-deficiency^18,19^. To progress from an animal model to a human-based model, induced pluripotent stem cells (iPSCs) were derived from AHDS patients and differentiated into brain microvascular endothelial-like cells, which showed MCT8-dependent transport of THs across the human BBB^16,20^. However, MCT8 is not restricted to the brain endothelium, and it also effects TH transport across neural cell plasma membranes^21^.

Gene therapy is a promising approach to treat monogenic genetic disorders. Spinal muscular atrophy (SMA) type 1 patients, carrying deleterious mutations in the *SMN1* gene, were treated by a single intravenous infusion of adeno-associated virus serotype 9 (AAV9) containing DNA coding for SMN, resulting in improved survival, as well as achievement of motor milestones and motor functions^22,23^, with subsequent FDA approval of the gene therapy product Zolgensma. Additional examples for beneficial gene therapy approach for monogenic disorders have been reported recently^24–26^.

While intracerebroventricular (ICV)-delivery directly targets the brain ventricles, thereby circumventing the blood brain barrier (BBB), intravenous (IV)-delivery offers systemic delivery, transducing primarily tissues outside the CNS including blood vessels. Importantly, it has been shown that AAV9 IV-delivery can cross the BBB and efficiently infect CNS cells^27^. In a recent proof of concept study, an AAV9-MCT8 construct was delivered by ICV or IV injections into neonatal *Mct8-*KO mice^28^, with an increase in brain TH signaling upon IV, but not ICV, delivery. However, since the *Mct8*-KO mice do not display neurological impairments, it was unclear whether this approach results in rescue of the neurological symptoms.

Here, we tested the potential of AAV9-MCT8 delivered IV to postnatal day 30 (P30) juvenile dKO (*mct8*^*-/y*^; *oatp1c1*^*-/-*^) mice. This approach was espoused as the diagnosis of MCT8 deficiency is usually made in childhood. Treatment resulted in expression of human MCT8 in the liver and throughout the brain, improved locomotor and cognitive behavior as well as complete and partial rescue of T_3_ content and associated gene expression in different areas of the brain. These preclinical findings provide the foundation for a future clinical trial using gene therapy for AHDS patients.

## Materials and Methods

### Experimental animals

All procedures were approved by Cedars-Sinai Medical Center’s Institutional Animal Care and Use Committee (IACUC #009128). *mct8*^*-*^*/*^*-*^; *oatp1c1*^*-*^*/*^*-*^ females and *mct8*^*-/y*^; *oatp1c1*^*-/-*^ males with C57BL/6 background were paired to generate dKO pups. Wildtype (WT) C57BL/6 were used as controls. Genders were identified at P1 by visual assessment and only males were selected for all treatments. Genotyping of *mct8* and *oatp1c1* were performed as previously described^17^. AAV9-MCT8 was administered to P30 juvenile dKO mice and controls by tail vein injection containing 50 × 10^10^ viral particles vp/g in a volume of 20 µl/g. Behavioral as well as biochemical and molecular measurements performed on tissues and serum were all performed and analyzed double blinded without the knowledge to which group the mice belonged. This was unblinded when results were assembled and for the statistical analysis.

### Virus

AAV9 was produced as previously described^29^. The short isoform of human MCT8 cDNA encoding a protein of 539 amino acids construct was previously cloned into the polylinkers of dsAAV CB MCS^28^, resulting in the AAV9-MCT8 vector. The virus was produced in HEK293 cells by transient transfection followed by purification using gradient density steps, dialyzed against phosphate-buffered saline (PBS) with 0.001% Pluronic-F68 to prevent virus aggregation, and was stored at 4°C. All vector preparations were titrated by quantitative real-time polymerase chain reaction (qRT-PCR) using Taq-Man technology.

### Mouse tissue collection

Mice were fully anesthetized at P140 with isoflurane, perfused with phosphate buffered saline (PBS) and then the brain and liver were collected, flash frozen on dry ice, and kept at -80°C until further processing. The hippocampus, thalamus, cerebral parietal cortex, choroid plexus, pituitary were subsequently dissected out for further analysis.

### Measurements of T_3_, T_4_, rT_3_ and TSH in serum and tissues

The tissue extraction procedure for T_3_ content, the measurements of serum TSH, T_4_, T_3_ and rT_3_ and tissue T_3_ content has previously been described in detail^30^. Briefly, Serum TSH was measured in 40 µl of serum using a double-antibody precipitation radioimmunoassay (RIA). The limit of detection was 10 mU/L.

For T_3_ measurement, 30 µl serum or tissue homogenates containing 1-2 mg of protein in PBS buffer (PH 7.4) were first extracted with chloroform-methanol (2:1, volume ratio). The remaining chloroform-phase was extracted with 0.05% CaCl_2_-MeOH-chloroform (48:49:3, volume ratio). The aqueous phase from both steps were collected, dried by vacuum evaporation and dissolved in T_3_ RIA buffer. The blanks and control samples were extracted at same time to correct the background and inter-assay variation. Recovery rate was 70-80%. T_3_ measurement was performed using a specific antibody against T_3_^31^. The sensitivity was 1 pg T_3_ per tube.

Total T_4_ was measured in 25 µl serum by coated-tube T_4_ monoclonal RIA from MP Biomedicals (Irvine, CA), the limit of detection was 0.5 µg/dL. Total rT_3_ was measured in 35 µl serum by RIA kit from (DIAsource ImmunoAssays S.A.). The limit of detection was 2.5 ng/dL.

### Extraction of tissue mRNA and qRT-PCR

RNA was extracted and used for measurement of specific mRNAs by qRT-PCR as previously described^32^. Amplification of the housekeeping gene Polymerase (RNA) II polypeptide A (Polr2a) was used as internal control. Primer sequences are provided in supplemental table 1.

### Behavioral tests

All behavioral assessments were performed at P120-P140 blinded following previously described protocols^17,21,33^ and are briefly described below.

### Rotarod

Mice were tested for motor coordination activity using an accelerating rotarod. The settings were changed to an acceleration of the rod rotation from 5 to 50 RPM within the maximum testing period of 300 s. The animals were analyzed for five consecutive days with three consecutive trials per day to determine the average time and distance the mice remained on the rotarod.

### Open field

Mice were placed in an open-topped, clear plexiglass chamber and left undisturbed for 30 min. Activity (includes horizontal locomotion and vertical rearing), speed and distance were tracked by disturbances in a grid of photobeams in the chamber, which were recorded using tracking software (Noldus, Leesburg, VA) and analyzed at 5-min intervals.

### Gait analysis

Mice were pretrained to run on a piece of filter paper. At P120 days, nontoxic ink was applied to the hind paws of the mice, and each animal walked on the filter paper at least 3 times. Paw print traces were analyzed using ImageJ (NIH) software with regard to the stride length and the hind paw angle.

### Barnes maze

As described previously^33^, mice were placed under bright light in the center of an elevated circular platform with 20 holes around the perimeter, one of which provided access to a small dark chamber (escape box). Colored shapes were placed around the maze to serve as visual cues. Mice were trained to find the escape box 3 times per day for 4 days (training phase, days 1–4). After a 2-day break, mice were tested on the maze (day 7) and the number of incorrect entries and time to find the escape box were recorded. The following day, the escape box was moved to the opposite side of the maze (reversal phase) and the mice were tested again for 2 consecutive days (days 8– 9).

### Y-maze

Mice were placed in an opaque, Y-shaped maze (San Diego Instruments, San Diego, CA) and spontaneous alternation of the mice between the three arms of the maze was recorded for 5 min.

### Statistical analysis

Results were unblinded for statistical analysis. All data are presented as mean ± SEM. p < 0.05 was considered statistically significant. *P < 0.05, **P < 0.01, ***P < 0.001, ****P < 0.0001. All scatter plots were first tested for their normal distribution using Kolmogorov-Smirnov test, with the Dallal-Wilkinson-Lilliefor corrected P value. Data sets that were normally distributed were tested using one-way ANOVA with Tukey’s post-hoc test for multiple comparisons. Data sets that were not normally distributed were compared using Mann-Whitney non-parametric test for independent samples. Behavior data collected over time was analyzed with mixed model regression with random intercept and the fixed-factors of time (as a continuous variable), treatment group, and the interaction term of group with time. To compare learning curves between groups the β coefficients were compared and differences were considered significant at the step-down Bonferroni-corrected alpha level of <0.05. Residuals were inspected to confirm the fit of the modeling. Data analysis was performed with GraphPad Prism v8.0.0 and SAS Enterprise Guide v8.2.

## Results

### IV delivery of AAV9-MCT8 at P30 improves locomotor performance of dKO mice

Given that patients with AHDS are often diagnosed during childhood, treatment feasibility should optimally be tested at a juvenile stage. Clinical applicability and previous experience^28^ indicate that IV delivery is more feasible compared to ICV delivery, as IV is a simpler route and would target not only the brain but other body regions that express MCT8 such as the liver. Therefore we tested IV delivery of 50 × 10^10^ vp/g of AAV9-MCT8 to peri-pubertal P30 dKO mice (Figure 1A). indicate

**Figure 1.**
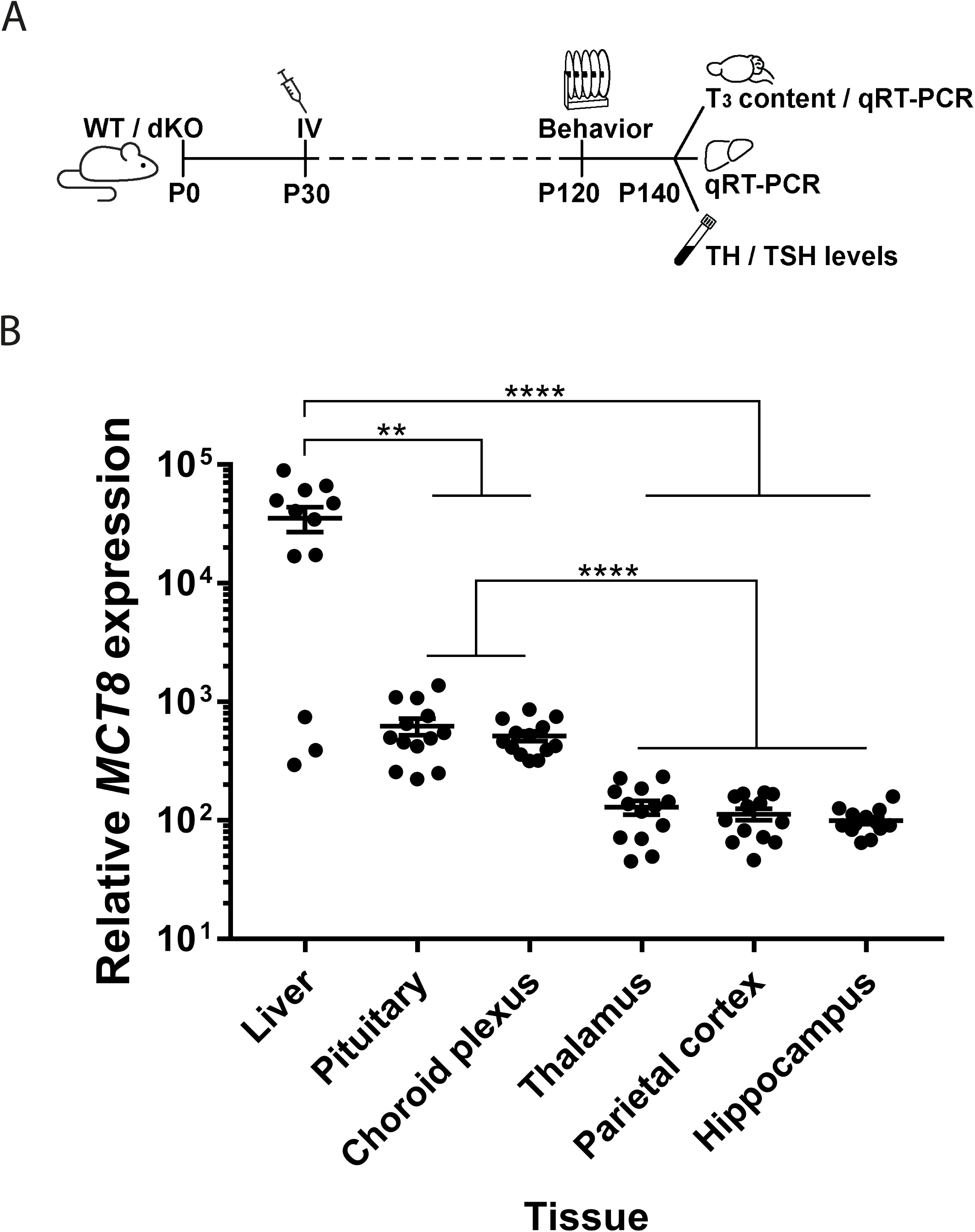
Study design and human *MCT8* expression in the liver and brain regions after IV AAV9-MCT8 injection of P30 dKO mice. **(A**) Schematic of experimental design. dKO mice were treated at postnatal day 30 (P30) by tail vain (IV) delivery of AAV9-MCT8 at a dose of 50 × 10^10^ vp/g. (**B**) Quantification of *MCT8* mRNA levels by qRT-PCR showed MCT8 re-expression in the liver and different brain regions of dKO treated animals. One-way ANOVA with Tukey’s multiple comparisons. The data are presented as mean, error bars represent SEM. (*P < 0.05, **P < 0.01, ***P < 0.001, ****P < 0.0001).

In order to confirm that the viral delivery resulted in expression of human *MCT8*, liver and brain were collected at P140. As expected, human *MCT8* mRNA expression was not detected in WT or dKO untreated mice (data not shown). *MCT8* expression was observed in the liver and various brain regions of AAV9-MCT8 IV-treated dKO mice, with the highest levels seen in the liver. Pituitary and choroid plexus had higher *MCT8* levels compared to the BBB-protected regions of the thalamus, parietal cortex and hippocampus (Figure 1B).

Mice underwent various behavioral analyses to assess locomotor function at P120. Assessment by a rotarod test showed that untreated dKO mice had an overall reduced latency to fall and decreased learning curve compared to the WT group (Figure 2A). Following IV delivery of AAV9-MCT8, dKO mice showed a significant increase in the latency to fall and an increased learning curve, indicating that treatment improved locomotor performance and potentially cognitive function. Overall locomotor activity assessed by an open field test as well as rearing behavior were both significantly higher in untreated dKO mice compared to WT mice, and were reduced to WT levels following treatment (Figure 2B and 2C). Finally, hind paw analysis showed that there was a partial improvement in the stride length and a complete recovery in the hind paw angle (Figure 2D and 2E). Collectively the data demonstrate that AAV9-MCT8 treatment at P30 provided some recovery of locomotion in dKO mice.

**Figure 2.**
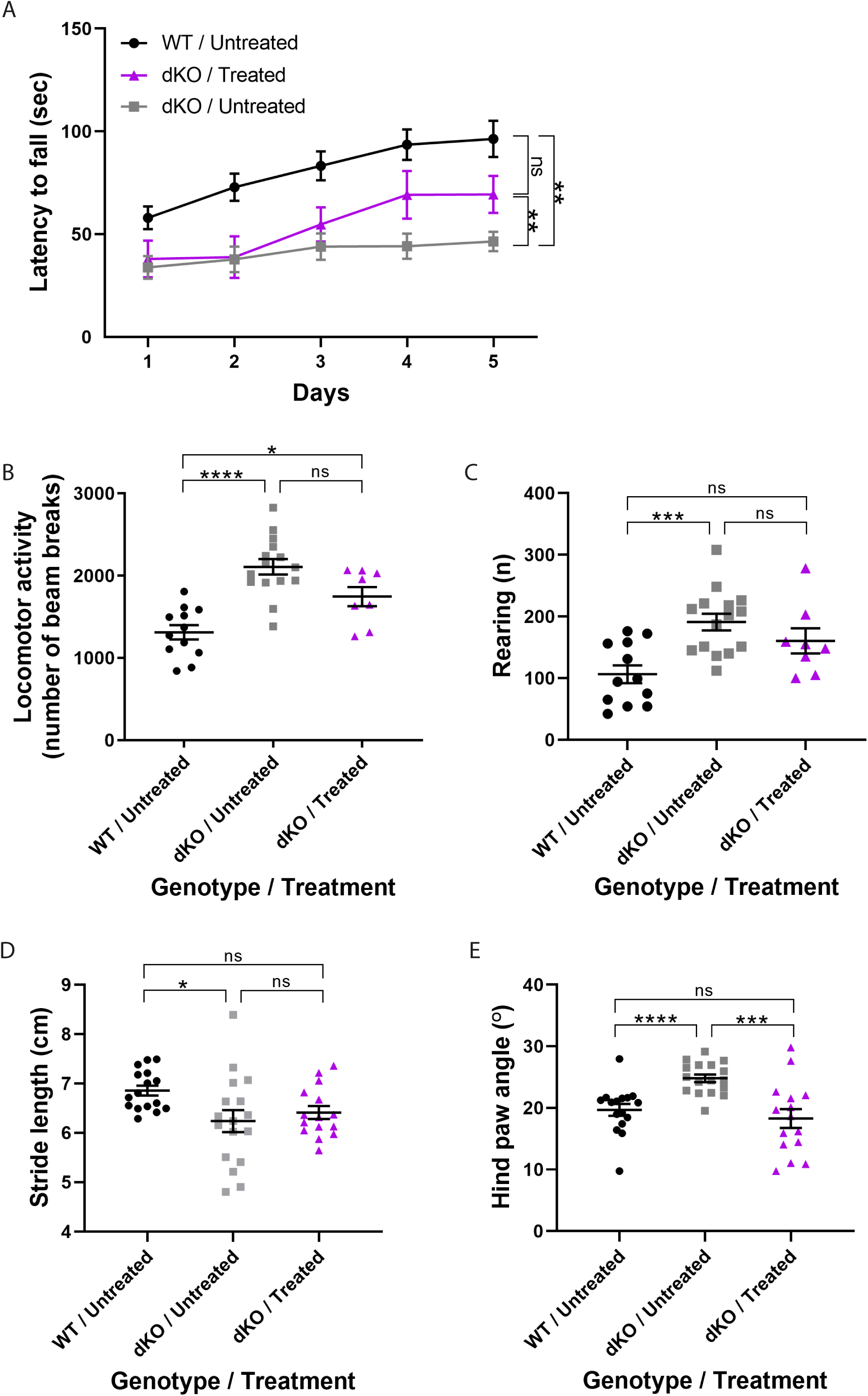
Locomotor performance is improved in dKO mice treated at P30. (**A**) Locomotor deficiencies were monitored by a rotarod test. Data was analyzed with mixed model regression with random intercept and the fixed-factors of time, group, and the interaction term of group with time. To compare learning curves between groups the β coefficients were compared. WT untreated mice (black line, n=16), treated dKO mice (purple line, n=13), untreated dKO mice (grey line, n=24). Open field test included (**B**) horizontal locomotion and (**C**) vertical rearing. (**D and E**) Paw print assessment at P120 to determine (**D**) Stride length and For B-D One-way ANOVA with Tukey’s multiple comparisons was used. (**E**) Hind paw angle. For E, Mann-Whitney non-parametric test for independent samples was used. The data are presented as mean, error bars represent SEM. (*P < 0.05, **P < 0.01, ***P < 0.001, ****P < 0.0001).

### IV delivery of AAV9-MCT8 at P30 improves cognitive performance in dKO mice

Learning, spatial memory and memory recall were assessed using a Barnes maze^33^. During the 4-day training period, treatment did not significantly improve the learning curve (Figure 3A). Following a 2-day break, untreated dKO mice required higher latency than untreated WT and treated dKO mice (Figure 3B). The position of the escape hole was then moved during two days of training during which no significant differences were observed between all groups (Figure 3C). These data suggest that the AAV9-MCT8 IV treatment of dKO mice at P30 resulted in a partial rescue in the learning and recall ability. A spontaneous alternation maze (Y-maze) test was used to further examine spatial memory. Untreated dKO mice demonstrated a significant decrease in the percent of spontaneous alterations compared to untreated WT mice, which was less significant in the AAV9-MCT8-treated dKO mice (Figure 3D). Collectively, these results suggest that IV-treatment of P30 dKO mice with AAV9-MCT8 improves cognitive performance as well as restores some spatial and learning memory.

**Figure 3.**
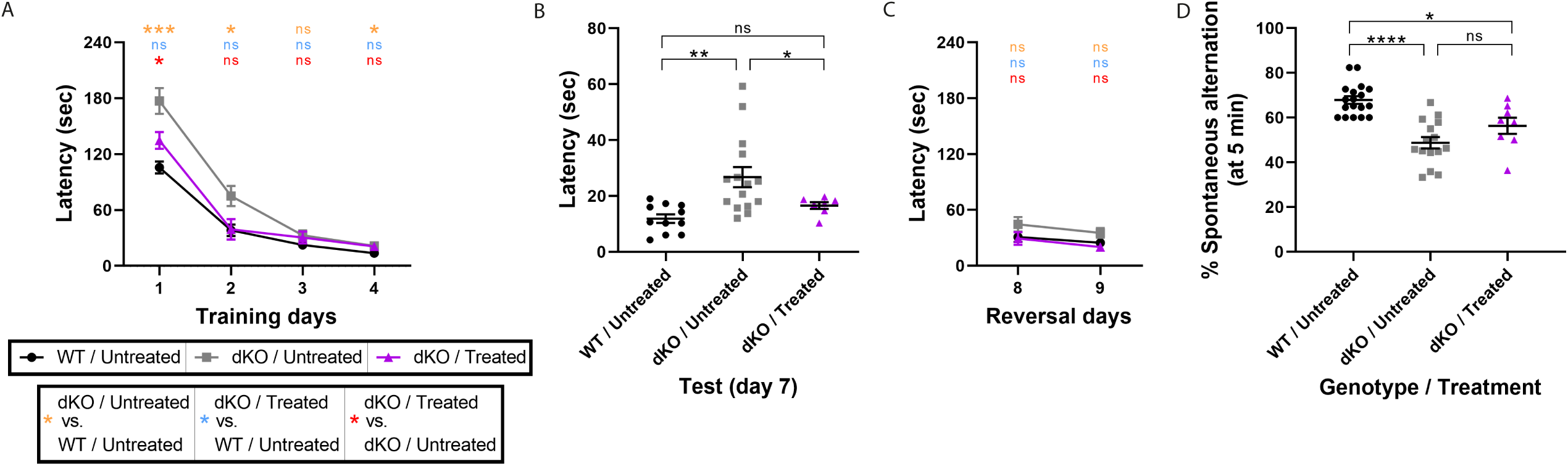
Improved cognitive performance in dKO mice treated at P30. The cognitive-related behavioral performance was assessed at P140. (**A-C**) In a Barnes Maze test, the ability of mice to discover and then recall the location of an escape hole was evaluated during the learning phase (**A**, Training, day 1–4), after a 2-day break (**B**, Test, day 7), and following re-positioning of the escape hole (**C**, Reversal day 8 and 9). The latency to successful location of the escape hole was recorded. Data was analyzed with mixed model regression with random intercept and the fixed-factors of time, group, and the interaction term of group with time. To compare learning curves between groups the β coefficients were compared. (**D**) Spontaneous alternation between the arms of a Y-maze was assessed over a 5-min period. One-way ANOVA with Tukey’s multiple comparisons. The data are presented as mean, error bars represent SEM. (*P < 0.05, **P < 0.01, ***P < 0.001, ****P < 0.0001). All analyses were double-blind.

### IV delivery of AAV9-MCT8 at P30 partially restores brain T_3_-content in dKO mice

In order to assess effects on pathophysiology, the serum, liver and brain were collected at P140 (Figure 1A). Examining the T_3_ brain content showed that T_3_ levels in the thalamus (Figure 4A) and hippocampus (Figure 4B) of treated dKO mice were fully normalized reaching WT levels. A significant increase in brain T_3_ content was also observed in the parietal cortex (Figure 4C).

**Figure 4.**
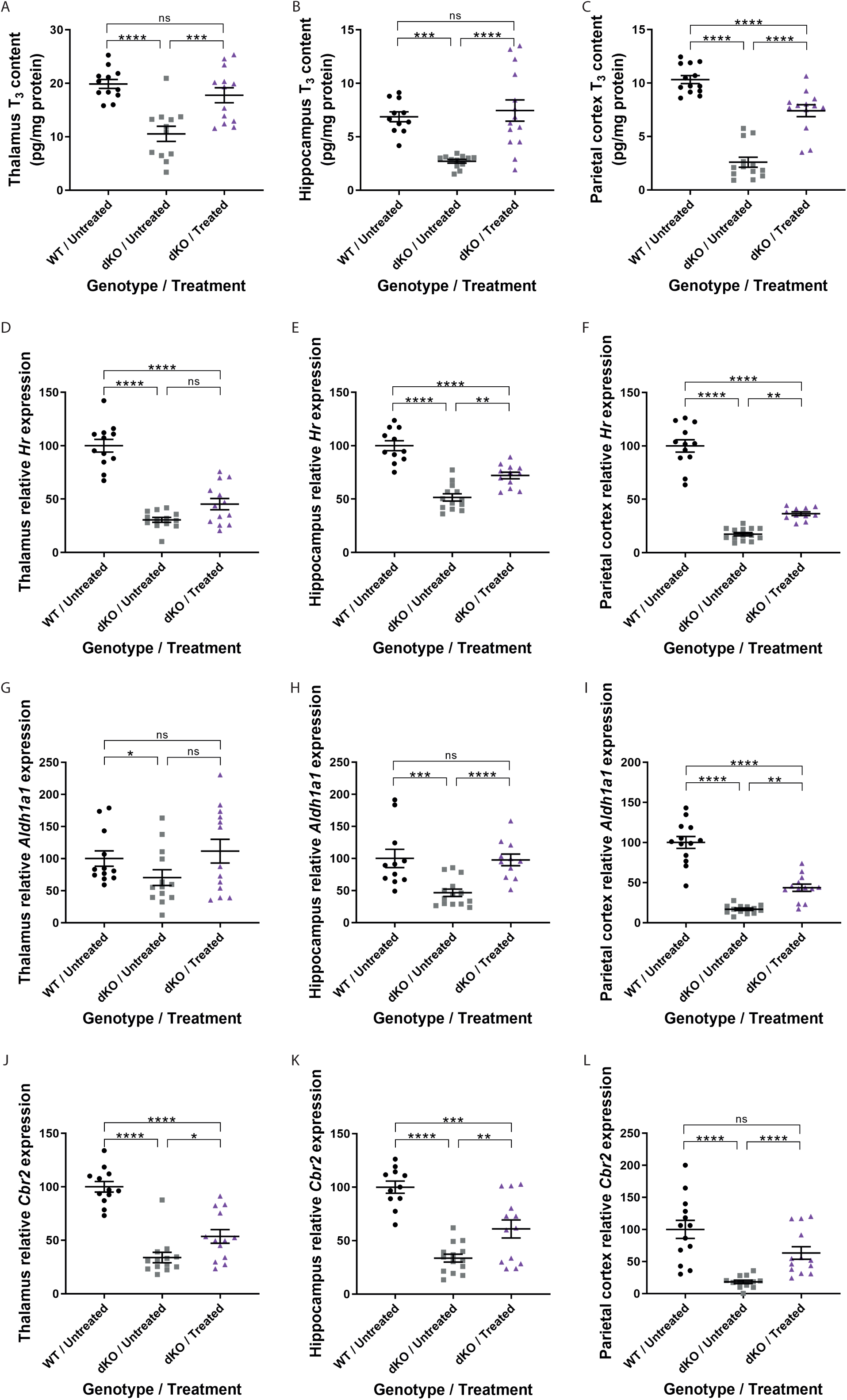
Brain T_3_ content and T_3_-induced gene expression in dKO mice treated at P30. T_3_ content measured in the (**A**) thalamus (**B**) hippocampus and (**C**) parietal cortex. T_3_-induced genes were examined by qRT-PCR. *Hairless* (*Hr*) expression was measured in the (**D**) thalamus (**E**) hippocampus and (**F**) parietal cortex. Expression of *Aldehyde dehydrogenase family 1, subfamily A1* (*Aldh1a1*) was measured in the (**G**) thalamus (**H**) hippocampus and (**I**) parietal cortex. Expression of *Carbonyl reductase 2* (*Cbr2*) was measured in the (**J**) thalamus (**K**) hippocampus and (**L**) parietal cortex. The data are presented as mean, error bars represent SEM. For C, G, H, J and L Mann-Whitney non-parametric test for independent samples was used. For A, B, D, E, F, I and K One-way ANOVA with Tukey’s multiple comparisons was used. The data are presented as mean, error bars represent SEM. (*P < 0.05, **P < 0.01, ***P < 0.001, ****P < 0.0001).

### IV delivery of AAV9-MCT8 at P30 corrects T_3_-inducible gene expression

To assess a T_3_-effect in the different brain regions, we next studied T_3_-inducible gene expression by qRT-PCR. *Hairless* (*Hr*) levels were slightly increased although not significantly in the thalamus of treated dKO mice compared to untreated dKO mice (Figure 4D) whereas levels were significantly improved in both the hippocampus (Figure 4E) and parietal cortex (Figure 4F) of treated dKO mice. *Aldehyde dehydrogenase 1 family member a1* (*Aldh1a1*) levels were not significantly different from untreated WT or dKO in the thalamus in response to treatment (Figure 4G), but were fully rescued in the hippocampus (Figure 4H) and significantly improved in the parietal cortex (Figure 4I). Finally, *Carbonyl reductase 2* (*Cbr2*) levels were also significantly improved in treated dKO compared to untreated dKO mice in the thalamus and hippocampus and restored in the parietal cortex (Figure 4J-L). These results suggest that IV delivery of AAV9-MCT8 to P30 dKO mice can restore and maintain long-term brain T_3_ content and T_3_-inducible gene expression.

### IV delivery of AAV9-MCT8 at P30 partially rescues phenotypes in the liver and serum of dKO mice

The significantly higher level of *MCT8* expression in the liver (Figure 1B) is expected with IV delivery, which can easily penetrate the blood vessels of the liver compared to the BBB capillaries. In contrast to the brain TH deficiency, patients with AHDS experience TH excess in peripheral tissues caused by the high serum T_3_ levels. We therefore measured the T_3_ effect in the liver by qRT-PCR of T_3_-inducible genes. The expression levels of *deiodinase 1* (*Dio1*), *malic enzyme 1* (*Me1*) and *uncoupling protein 2* (*Ucp2*) (Figure 5A-C) were all significantly elevated in the dKO untreated group compared to their WT littermates, confirming the effect of TH excess in liver. AAV9-MCT8 delivery led to a significant decrease in *Dio1* mRNA levels in treated dKO mice compared to untreated dKO mice. However, there was not complete correction of TH excess as AAV9-MCT8 treatment did not significantly reduce *Me1* and *Ucp2* levels.

**Figure 5.**
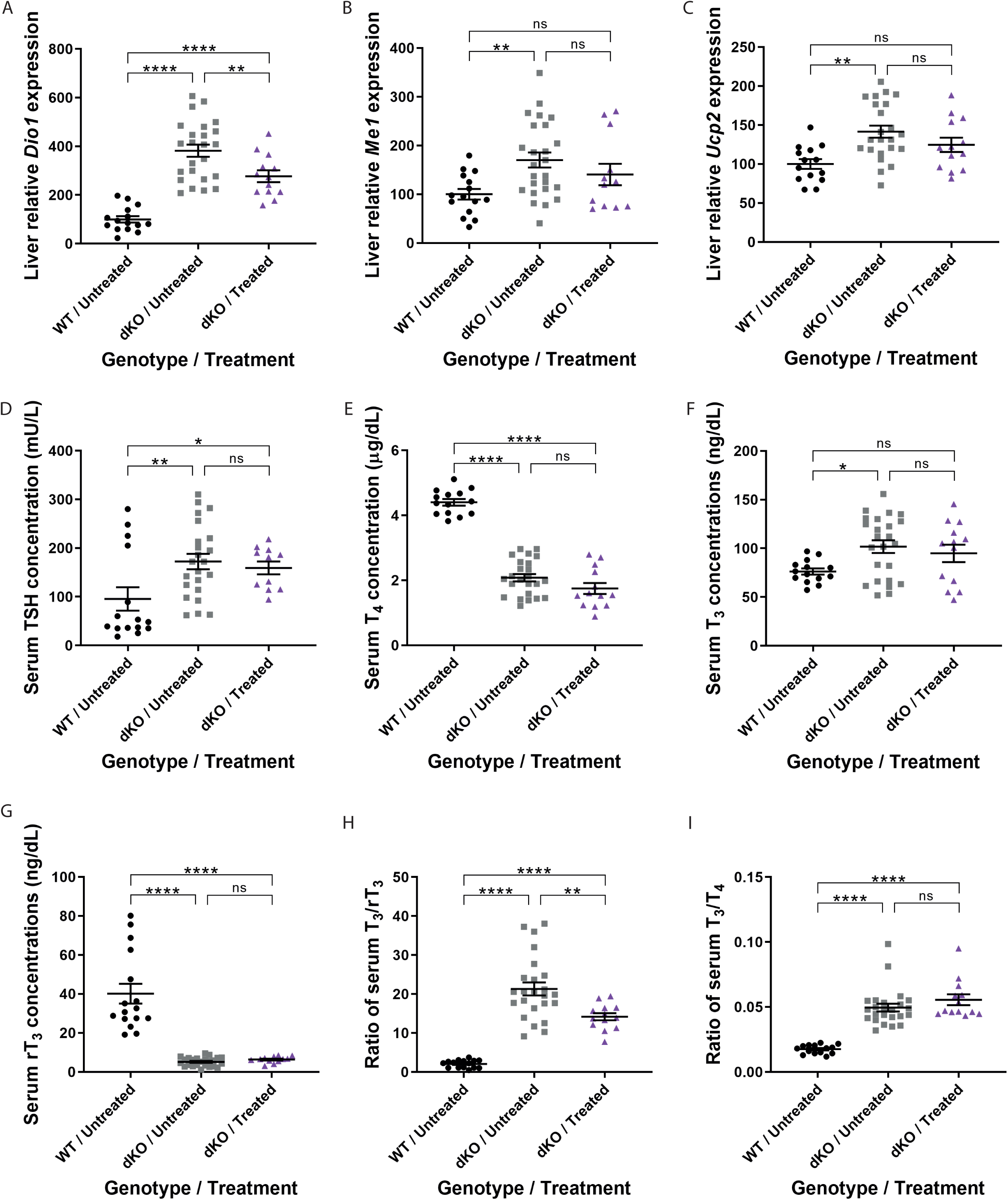
Liver T_3_-induced gene expression and serum TSH and TH concentrations in dKO mice treated at P30. Livers tissue was obtained for qRT-PCR analysis of T_3_-induced gene expression including (**A**) *iodothyronine deiodinase 1* (*dio1*), (**B**) *malic enzyme 1*(*Me1*) and (**C**) *uncoupling protein 2* (*Ucp2*). Concentrations of hormones in serum are shown including (**D**) TSH (**E**) T_4_, (**F**) T_3_, (**G**) and reverse T_3_ (rT_3_). Ratios of (**H**) T_3_/rT_3_ and (**I**) T_3_/T_4_ were calculated. Data are presented as mean, error bars represent SEM. For D, G and I Mann-Whitney non-parametric test for independent samples was used. For A-C, E, F and H One-way ANOVA with Tukey’s multiple comparisons was used. The data are presented as mean, error bars represent SEM. (*P < 0.05, **P < 0.01, ***P < 0.001, ****P < 0.0001).

Patients with AHDS have abnormal serum TH levels, including elevated T_3_, low rT_3_ and T_4_ with normal or slightly elevated TSH, resulting in low T_3_/T_4_ and T_3_/rT_3_ ratios. The increased liver deiodinase 1 enzymatic activity is one of the mechanisms responsible for these serum thyroid tests and a decrease in its expression is needed to ameliorate this phenotype^34^. In order to test the effect of IV delivery of AAV9-MCT8 at P30 on the serum TH phenotype in dKO mice, blood was collected from P140 mice and serum TSH and TH levels were quantified. Serum levels of TSH, T_4_, T_3_, and rT_3_ as well as the T_3_/rT_3_ and the T_3_/T_4_ ratios were all significantly altered in dKO mice compared to their WT littermates (Figure 5D-I). While AAV9-MCT8 treatment did not significantly alter serum levels of TSH, T_4_, T_3_, rT_3_ and T_3_/T_4_ ratio, the combination of slight reduction in T_3_ and increase in rT_3_ resulted in a significant decrease of the T_3_/rT_3_ ratio in agreement with the observed attenuation of *Dio1* expression. This indicates that AAV9-MCT8 delivery at a juvenile stage can partially improve the abnormalities in serum. These can be easily corrected with thyroid hormone analogues, which fail to improve the more serious consequences, namely, the neuropsychomotor defects of AHDS^35–37^.

## Discussion

AHDS is a severe X-linked psychomotor disability caused by inactivating mutations in the gene encoding MCT8^7,8^. In addition to the neuro-psychomotor phenotype, MCT8-deficient patients suffer from TH excess in peripheral tissues. Thus, an effective therapeutic strategy should account for deficient transport of THs across brain barriers and neural plasma membranes as well as the excess of TH in peripheral tissues. Being a rare disorder, AHDS is often misdiagnosed resulting in later identification of the disease. Moreover, there are currently about 300 diagnosed cases, which mostly involve older children^11,38^. Thus, it is important to develop therapeutics that are effective at juvenile ages. We therefore, tested the effect of tail vein IV delivery at P30, which is peri-pubertal when pathophysiological symptoms are apparent in both dKO mice and patients^17,39,40^.

There are several current treatment strategies that focus on TH analogs. These thyromimetic compounds are required to activate TH-induced transcriptional pathways through activation of thyroid nuclear receptors, and need to penetrate plasma membranes independent of MCT8. In animal models, the TH analogs DITPA^35,41^, TRIAC^18^, TETRAC^19^ and Sobetirome^42^ were able to restore some of the peripheral and central abnormalities, however, their effects on neurological symptoms remained limited or unknown due to the use of mice that were not neurologically affected. In patients with AHDS, DITPA^36^ and TRIAC^37^ reduced the high serum T_3_ concentration, however, there was no evidence for improvement in neurological symptoms.

Chemical and pharmaceutical chaperons have also been suggested as an alternative approach. These chaperons can restore the ability of some *MCT8* gene mutations to transport TH across plasma membranes in animal models^43,44^, however, to date their clinical effect has not been assessed. Moreover, this approach is limited only to some missense mutations, and is therefore not applicable to the majority of patients.

Recently, gene replacement therapy using adeno-associated viral serotype 9 (scAAV9) has emerged as a promising approach to treat monogenic developmental neurological disorders^23,25,26,45–47^. Unlike TH-analogs, restoration of a functional MCT8 has the potential to resolve TH transport, as well as unidentified MCT8 roles. Further, gene replacement can potentially target all mutations and patients. A recent study tested the delivery of AAV9-MCT8 in *Mct8*-KO mice^28^, demonstrating that IV, but not ICV, injection at a neonatal stage increases brain T_3_ content and T_3_-induced gene expression. Since MCT8 has a crucial role in the delivery of THs across the BBB^16,17,20,21,48,49^, IV delivery, targeting endothelial cells of the BBB, may provide a more suitable approach to potentially treat MCT8-deficient patients. However, as *Mct8*-KO mice fail to display disease-relevant neurological phenotypes, the rescue of neurological symptoms could not be assessed. Additionally, the serum and peripheral tissue phenotypes characteristic of AHDS were not evaluated. Therefore, here, we tested whether IV delivery of AAV9-MCT8 can recue both the psychomotor and systemic phenotype using the well-established dKO (*mct8*^*-/y*^; *oatp1c1*^*-/-*^) mouse model that has several AHDS-relevant phenotypes^17,50^.

MCT8 is ubiquitously expressed and is prominently localized in the thyroid, liver, kidneys and CNS^6,12,13^. In the current study we show that IV-administration of AAV9-MCT8 to P30 dKO mice led to long-term *MCT8* expression in the liver and CNS. Liver *MCT8* expression was significantly higher than in the brain and was accompanied by a reduction in *Dio1* expression, a TH-regulated enzyme that generates T_3_ from T_4_ and is responsible in part for the T_3_ excess in serum^34^. This effect on *Dio1* expression resulted in a partial amelioration in serum TH parameters in particular the T_3_/rT_3_ ratio. These results suggest a beneficial effect of systemic (IV) delivery of MCT8 on the peripheral tissues. Additional mechanisms contribute to the characteristic serum thyroid tests of AHDS, including decreased thyroidal secretion and altered negative feedback to the hypothalamus, which is BBB-protected, and pituitary, located outside of BBB, and thus have different TH availability^51–53^. Another layer of complexity in the circulating and intracellular TH levels is added by the multifaceted regulation of the three deiodinase enzymes that metabolize TH and are also controlled by TH either through direct genomic effect (D1 and D3) or post-translational (D2)^2^. Thus, to augment the partial rescue, additional TH-normalizing treatments should be considered in conjunction with gene therapy.

*MCT8* expression was observed in various regions of the brain, confirming the ability of AAV9 vectors to cross brain barriers and efficiently transduce brain cells^27^. Interestingly, higher expression was observed in brain regions that are not protected by the BBB such as the hypothalamus and pituitary^54,55^ compared to the thalamus, hippocampus and cortex. Strikingly, in the current study this long-term brain expression resulted in a nearly complete normalization of T_3_ brain concentrations and some associated gene expressions. Treated animals also showed an improvement in the learning curve using the rotarod test, suggesting that treatment may have beneficial effects on cognitive and motor functions. Assessment with cognitive tests showed that treated dKO mice displayed mild improved cognition in the Barnes maze test. Exploratory behavior, learning and memory are thought to originate in the hippocampus^56^. The significant rescue of hippocampus-dependent learning and memory in response to treatment corresponded with the significant increase of T_3_ levels and T_3_-induced gene expression in the hippocampus. Collectively, these results indicate that IV delivery of MCT8 at P30 rescues both psychomotor and cognitive deficiency in dKO mice.

Overall, this study shows that IV administration of AAV9-MCT8 to dKO mice provides substantial rescue of molecular and biochemical parameters in the brain, and an amelioration of the TH excess effect in peripheral tissues. In addition, this treatment improves locomotor and behavioral performance. These findings offer the preclinical foundation for future clinical examination of AAV-based MCT8 gene therapy in patients with AHDS.

## Acknowledgements

The authors would like to thank Dr. Soshana Svendsen for critical writing and editing. Biostatistical support was received from the Center for Research Support (“HALEV”), Faculty of Health Sciences, Ben Gurion University of the Negev and by the Samuel Oschin Comprehensive Cancer Institute at Cedars-Sinai. This work was supported by the Sherman Family Foundation, the Board of Governors Regenerative Medicine Institute at Cedars-Sinai Medical Center to CNS, the Israel Science Foundation grant 1621/18 and the Ministry of Science and Technology (MoST), Israel grant 3-15647 to GDV. This work was supported in part by grant DK15070 from the National Institutes of Health (USA) to SR.

## Author contribution

GDV, SR and CNS provided the conceptual framework for the study and designed the experiments. XHL performed the brain dissections, qRT-PCR analyses and the TH measurements. PA treated the animals. OS bred the colonies and collected tissues. OS and JPV performed behavioral analyses. RO analyzed the data. MS and CB performed the statistical analyses. SL, BK and KM provided the virus. AMD generated the human MCT8 insert used in the construction of the virus and provided intellectual input. HH provided the dKO mice and provided intellectual input. GDV and CNS wrote the manuscript.

## References

1. Yen, P. M. Physiological and molecular basis of Thyroid hormone action. Physiological Reviews vol. 81 (2001).

2. Gereben, B. et al. Cellular and molecular basis of deiodinase-regulated thyroid hormone signaling. Endocrine Reviews vol. 29 (2008).

3. Harvey, C. B. & Williams, G. R. Mechanism of thyroid hormone action. Thyroid vol. 12 (2002).

4. Groeneweg, S., Van Geest, F. S., Peeters, R. P., Heuer, H. & Visser, W. E. Thyroid Hormone Transporters. Endocrine Reviews vol. 41 (2019).

5. Friesema, E. C. H. et al. Identification of monocarboxylate transporter 8 as a specific thyroid hormone transporter. J. Biol. Chem. (2003) doi:10.1074/jbc.M300909200.

6. Vatine, G. D. et al. Zebrafish as a model for monocarboxyl transporter 8-deficiency. J. Biol. Chem. (2013) doi:10.1074/jbc.M112.413831.

7. Dumitrescu, A. M., Liao, X. H., Best, T. B., Brockmann, K. & Refetoff, S. A Novel Syndrome Combining Thyroid and Neurological Abnormalities Is Associated with Mutations in a Monocarboxylate Transporter Gene. Am. J. Hum. Genet. (2004) doi:10.1086/380999.

8. Friesema, E. C. H. et al. Association between mutations in a thyroid hormone transporter and severe X-linked psychomotor retardation. Lancet (2004) doi:10.1016/S0140-6736(04)17226-7.

9. Schwartz, C. E. et al. Allan-Herndon-Dudley syndrome and the monocarboxylate transporter 8 (MCT8) gene. Am. J. Hum. Genet. (2005) doi:10.1086/431313.

10. Allan, W., Herndon, C. N. & Dudley, F. C. Some examples of the inheritance of mental deficiency: apparently sex-linked idiocy and microcephaly. Am. J. Ment. Defic. (1944).

11. Groeneweg, S. et al. Disease characteristics of MCT8 deficiency: an international, retrospective, multicentre cohort study. Lancet Diabetes Endocrinol. (2020) doi:10.1016/S2213-8587(20)30153-4.

12. Dumitrescu, A. M., Liao, X. H., Weiss, R. E., Millen, K. & Refetoff, S. Tissue-specific thyroid hormone deprivation and excess in monocarboxylate transporter (Mct) 8-deficient mice. Endocrinology (2006) doi:10.1210/en.2006-0390.

13. Trajkovic, M. et al. Abnormal thyroid hormone metabolism in mice lacking the monocarboxylate transporter 8. J. Clin. Invest. (2007) doi:10.1172/JCI28253.

14. Mayerl, S., Visser, T. J., Darras, V. M., Horn, S. & Heuer, H. Impact of Oatp1c1 deficiency on thyroid hormone metabolism and action in the mouse brain. Endocrinology (2012) doi:10.1210/en.2011-1633.

15. Roberts, L. M. et al. Expression of the thyroid hormone transporters monocarboxylate transporter-8 (SLC16A2) and organic ion transporter-14 (SLCO1C1) at the blood-brain barrier. Endocrinology (2008) doi:10.1210/en.2008-0378.

16. Vatine, G. D. et al. Modeling Psychomotor Retardation using iPSCs from MCT8-Deficient Patients Indicates a Prominent Role for the Blood-Brain Barrier. Cell Stem Cell (2017) doi:10.1016/j.stem.2017.04.002.

17. Mayerl, S. et al. Transporters MCT8 and OATP1C1 maintain murine brain thyroid hormone homeostasis. J. Clin. Invest. (2014) doi:10.1172/JCI70324.

18. Kersseboom, S. et al. In vitro and mouse studies support therapeutic utility of triiodothyroacetic acid in MCT8 deficiency. Mol. Endocrinol. (2015) doi:10.1210/me.0000-9999.

19. Horn, S. et al. Tetrac can replace thyroid hormone during brain development in mouse mutants deficient in the thyroid hormone transporter Mct8. Endocrinology (2013) doi:10.1210/en.2012-1628.

20. Vatine, G. D. et al. Human iPSC-Derived Blood-Brain Barrier Chips Enable Disease Modeling and Personalized Medicine Applications. Cell Stem Cell (2019) doi:10.1016/j.stem.2019.05.011.

21. Vatine, G. D. et al. Oligodendrocyte progenitor cell maturation is dependent on dual function of MCT8 in the transport of thyroid hormone across brain barriers and the plasma membrane. Glia 69, (2021).

22. Mendell, J. R. et al. Single-Dose Gene-Replacement Therapy for Spinal Muscular Atrophy. N. Engl. J. Med. 377, (2017).

23. Mendell, J. R. et al. Five-Year Extension Results of the Phase 1 START Trial of Onasemnogene Abeparvovec in Spinal Muscular Atrophy. JAMA Neurol. 78, (2021).

24. Corti, M. et al. Safety of Intradiaphragmatic Delivery of Adeno-Associated Virus-Mediated Alpha-Glucosidase (rAAV1-CMV-hGAA) Gene Therapy in Children Affected by Pompe Disease. Hum. Gene Ther. Clin. Dev. 28, (2017).

25. Mendell, J. R. et al. Assessment of Systemic Delivery of rAAVrh74.MHCK7.micro-dystrophin in Children with Duchenne Muscular Dystrophy: A Nonrandomized Controlled Trial. JAMA Neurol. 77, (2020).

26. Eichler, F. et al. Hematopoietic Stem-Cell Gene Therapy for Cerebral Adrenoleukodystrophy. N. Engl. J. Med. 377, (2017).

27. Foust, K. D. et al. Intravascular AAV9 preferentially targets neonatal neurons and adult astrocytes. Nat. Biotechnol. 27, (2009).

28. Iwayama, H. et al. Adeno Associated Virus 9-Based Gene Therapy Delivers a Functional Monocarboxylate Transporter 8, Improving Thyroid Hormone Availability to the Brain of Mct8-Deficient Mice. in Thyroid (2016). doi:10.1089/thy.2016.0060.

29. Foust, K. D. et al. Therapeutic AAV9-mediated suppression of mutant SOD1 slows disease progression and extends survival in models of inherited ALS. Mol. Ther. 21, (2013).

30. Ferrara, A. M. et al. Changes in thyroid status during perinatal development of MCT8-deficient male mice. Endocrinology 154, (2013).

31. Obregon, M. J., Pascual, A., De Escobar, G. M. & Del Rey, F. E. Pituitary and plasma thyrotropin, thyroxine, and triiodothyronine after hyperthyroidism. Endocrinology 104, (1979).

32. Fujisawa, H., Korwutthikulrangsri, M., Fu, J., Liao, X. H. & Dumitrescu, A. M. Role of the thyroid gland in expression of the thyroid phenotype of Sbp2-deficient mice. Endocrinol. (United States) 161, (2020).

33. Das, M. M. et al. Young bone marrow transplantation preserves learning and memory in old mice. Commun. Biol. 2, (2019).

34. Di Cosmo, C. et al. Mct8-deficient mice have increased energy expenditure and reduced fat mass that is abrogated by normalization of serum T3 levels. Endocrinology 154, (2013).

35. Di Cosmo, C., Liao, X. H., Dumitrescu, A. M., Weiss, R. E. & Refetoff, S. A thyroid hormone analog with reduced dependence on the monocarboxylate transporter 8 for tissue transport. Endocrinology 150, (2009).

36. Verge, C. F. et al. Diiodothyropropionic acid (DITPA) in the treatment of MCT8 deficiency. J. Clin. Endocrinol. Metab. 97, (2012).

37. Groeneweg, S. et al. Effectiveness and safety of the tri-iodothyronine analogue Triac in children and adults with MCT8 deficiency: an international, single-arm, open-label, phase 2 trial. Lancet Diabetes Endocrinol. 7, (2019).

38. van Geest, F. S., Groeneweg, S. & Visser, W. E. Monocarboxylate transporter 8 deficiency: update on clinical characteristics and treatment. Endocrine vol. 71 (2021).

39. Bárez-López, S. et al. Adult Mice Lacking Mct8 and Dio2 Proteins Present Alterations in Peripheral Thyroid Hormone Levels and Severe Brain and Motor Skill Impairments. Thyroid (2019) doi:10.1089/thy.2019.0068.

40. López-Espíndola, D. et al. Mutations of the thyroid hormone transporter MCT8 cause prenatal brain damage and persistent hypomyelination. J. Clin. Endocrinol. Metab. (2014) doi:10.1210/jc.2014-2162.

41. Ferrara, A. M. et al. The thyroid hormone analog DITPA ameliorates metabolic parameters of male mice with Mct8 deficiency. Endocrinology (2015) doi:10.1210/en.2015-1234.

42. Bárez-López, S., Hartley, M. D., Grijota-Martínez, C., Scanlan, T. S. & Guadaño-Ferraz, A. Sobetirome and its Amide Prodrug Sob-AM2 Exert Thyromimetic Actions in Mct8-Deficient Brain. Thyroid 28, (2018).

43. Braun, D. & Schweizer, U. The chemical chaperone phenylbutyrate rescues MCT8 mutations associated with milder phenotypes in patients with Allan-Herndon-Dudley syndrome. Endocrinology 158, (2017).

44. Braun, D. & Schweizer, U. Efficient activation of pathogenic Δphe501 mutation in monocarboxylate transporter 8 by chemical and pharmacological chaperones. Endocrinology 156, (2015).

45. Mendell, J. R. et al. Single-dose gene-replacement therapy for spinal muscular atrophy. N. Engl. J. Med. (2017) doi:10.1056/NEJMoa1706198.

46. White, K. A. et al. Intracranial delivery of AAV9 gene therapy partially prevents retinal degeneration and visual deficits in CLN6-Batten disease mice. Mol. Ther. -Methods Clin. Dev. 20, (2021).

47. Pearson, T. S. et al. Gene therapy for aromatic L-amino acid decarboxylase deficiency by MR-guided direct delivery of AAV2-AADC to midbrain dopaminergic neurons. Nat. Commun. 12, (2021).

48. Ceballos, A. et al. Importance of monocarboxylate transporter 8 for the blood-brain barrier-dependent availability of 3,5,3’ -triiodo-L-thyronine. Endocrinology (2009) doi:10.1210/en.2008-1616.

49. Zada, D., Tovin, A., Lerer-Goldshtein, T. & Appelbaum, L. Pharmacological treatment and BBB-targeted genetic therapy for MCT8-dependent hypomyelination in zebrafish. DMM Dis. Model. Mech. (2016) doi:10.1242/dmm.027227.

50. Mayerl, S. et al. Thyroid Hormone Transporters MCT8 and OATP1C1 Control Skeletal Muscle Regeneration. Stem Cell Reports 10, (2018).

51. Trajkovic-Arsic, M. et al. Consequences of monocarboxylate transporter 8 deficiency for renal transport and metabolism of thyroid hormones in mice. Endocrinology 151, (2010).

52. Di Cosmo, C. et al. Mice deficient in MCT8 reveal a mechanism regulating thyroid hormone secretion. J. Clin. Invest. 120, (2010).

53. Fliers, E., Unmehopa, U. A. & Alkemade, A. Functional neuroanatomy of thyroid hormone feedback in the human hypothalamus and pituitary gland. Molecular and Cellular Endocrinology vol. 251 (2006).

54. Anbalagan, S. et al. Pituicyte Cues Regulate the Development of Permeable Neuro-Vascular Interfaces. Dev. Cell 47, (2018).

55. Ben-Zvi, A. & Liebner, S. Developmental regulation of barrier-and non-barrier blood vessels in the CNS. J. Intern. Med. (2021) doi:10.1111/joim.13263.

56. Voss, J. L., Gonsalves, B. D., Federmeier, K. D., Tranel, D. & Cohen, N. J. Hippocampal brain-network coordination during volitional exploratory behavior enhances learning. Nat. Neurosci. 14, (2011).

